# Familiarity facilitates detection of angry expressions

**DOI:** 10.1101/458984

**Authors:** Vassiki Chauhan, Matteo Visconti di Oleggio Castello, Morgan Taylor, Maria Ida Gobbini

**Affiliations:** Department of Psychological and Brain Sciences, Dartmouth College, Hanover, NH, USA; Department of Medicina Specialistica, Diagnostica e Sperimentale (DIMES), Medical School, University of Bologna, Italy

## Abstract

Personal familiarity facilitates rapid and optimized detection of faces. In this study, we investigated whether familiarity associated with faces can also facilitate the detection of facial expressions. Models of face processing propose that face identity and face expression detection are mediated by distinct pathways. We used a visual search paradigm to assess if facial expressions of emotion (anger and happiness) were detected more rapidly when produced by familiar as compared to unfamiliar faces. We found that participants detected an angry expression 11% more accurately and 135 ms faster when produced by familiar as compared to unfamiliar faces while happy expressions were detected with equivalent accuracies and at equivalent speeds for familiar and unfamiliar faces. These results suggest that detectors in the visual system dedicated to processing features of angry expressions are optimized for familiar faces.

## Introduction

Faces provide a vast range of information about people that we rely on for social interactions (Zebrowitz & Montepare, 2008; Todorov et al., 2007; Gobbini & Haxby 2007; Sugiura et al., 2014; Natu & O’Toole, 2011; Ramon & Gobbini, 2018; Jack & Schyns, 2015). Personal familiarity plays a critical role in tuning the face processing system (Gobbini et al. 2004; Leibenluft et al. 2004; Gobbini & Haxby, 2007; Natu & O’Toole, 2011; Ramon & Gobbini, 2018). We have shown that familiar faces are detected more readily even with reduced attention and without conscious awareness (Gobbini et al. 2013), and that familiar faces can be detected around 130 ms faster in a visual search task (Visconti di Oleggio Castello et al. 2017). Furthermore, we have proposed that detection of identity is, at least in part, supported by identity related feature detectors in retinotopic visual cortex that are strengthened by familiarity (Visconti di Oleggio Castello et al. 2018). We also have shown that eye gaze direction is detected around 100 ms faster in personally familiar faces, as compared to the faces of strangers, in a visual search task. This latter result suggests that detectors for facial-gesture-related features also are tuned by personal familiarity.

Here, we test whether facial expression also is detected more efficiently in personally familiar faces as compared to the faces of strangers. We used a visual search task in which participants report the presence or absence of a happy or an angry face target among faces with neutral expressions. We hypothesized that target detection is faster when the expression is conveyed by a personally familiar face than by an unfamiliar face. Results support our hypothesis, suggesting that detectors for facial expressions are tuned specifically for how these features appear in personally familiar faces, thereby facilitating optimized performance in social interactions with personally familiar others.

## Methods

### Participants

15 graduate students (9 female, age: 25.6 ± 2.0) from Dartmouth College participated in the experiment. The sample size was chosen to be consistent with previous studies using the same paradigm (Tong & Nakayama, 1999; Visconti di Oleggio Castello, Guntupalli, 2014; Visconti di Oleggio Castello, 2017). All participants had normal or corrected to normal vision. Participants were recruited from the graduate students of the department of Psychological and Brain Sciences who were part of the program for at least one year. All participants provided written informed consent to participate in the experiment, and were monetarily compensated for their time. The study was approved by the Dartmouth Committee for the Protection of Human Subjects (Protocol 29780).

### Stimuli

The stimulus set consisted of 34 images. 24 images were of target identities, 12 of which were personally familiar to all of the participants (fellow graduate students) and 12 of which were unfamiliar controls, chosen to be visually similar to the familiar identities. 10 images were of distractor identities (5 male and 5 female). To ensure that the graduate students were not visually familiar with the unfamiliar targets and the distractors, images of unfamiliar identities were collected at the Massachusetts Institute of Technology. Three images per target identity were used — one image for angry, happy and neutral expressions. In this experiment, we used 4 familiar identities (2 male and 2 female) and 4 unfamiliar identities (2 male and 2 female) matched for gender, ethnicity and age to the familiar targets. All stimuli were collected with the same camera, placement, settings and lighting equipment to minimize differences due to image quality. They were cropped to be the same size (350 × 350 pixels) using custom code written in Matlab. To further reduce low level visual differences, the average pixel intensity of each image (ranging from 0 to 255) was set to 128 using the SHINE toolbox function *lumMatch* in Matlab (Willenbockel et al., 2010).

During the task, stimuli were presented in sets of two, four or six. These images were presented symmetrically, placed in a regular hexagon, centered on the fixation cross, such that the center of each image was always at 7° of visual angle from the fixation cross (Figure 1). Each image subtended 4° × 4° of visual angle. The positions of the images were chosen to be symmetric around the fixation for each set size.

**Figure 1:**
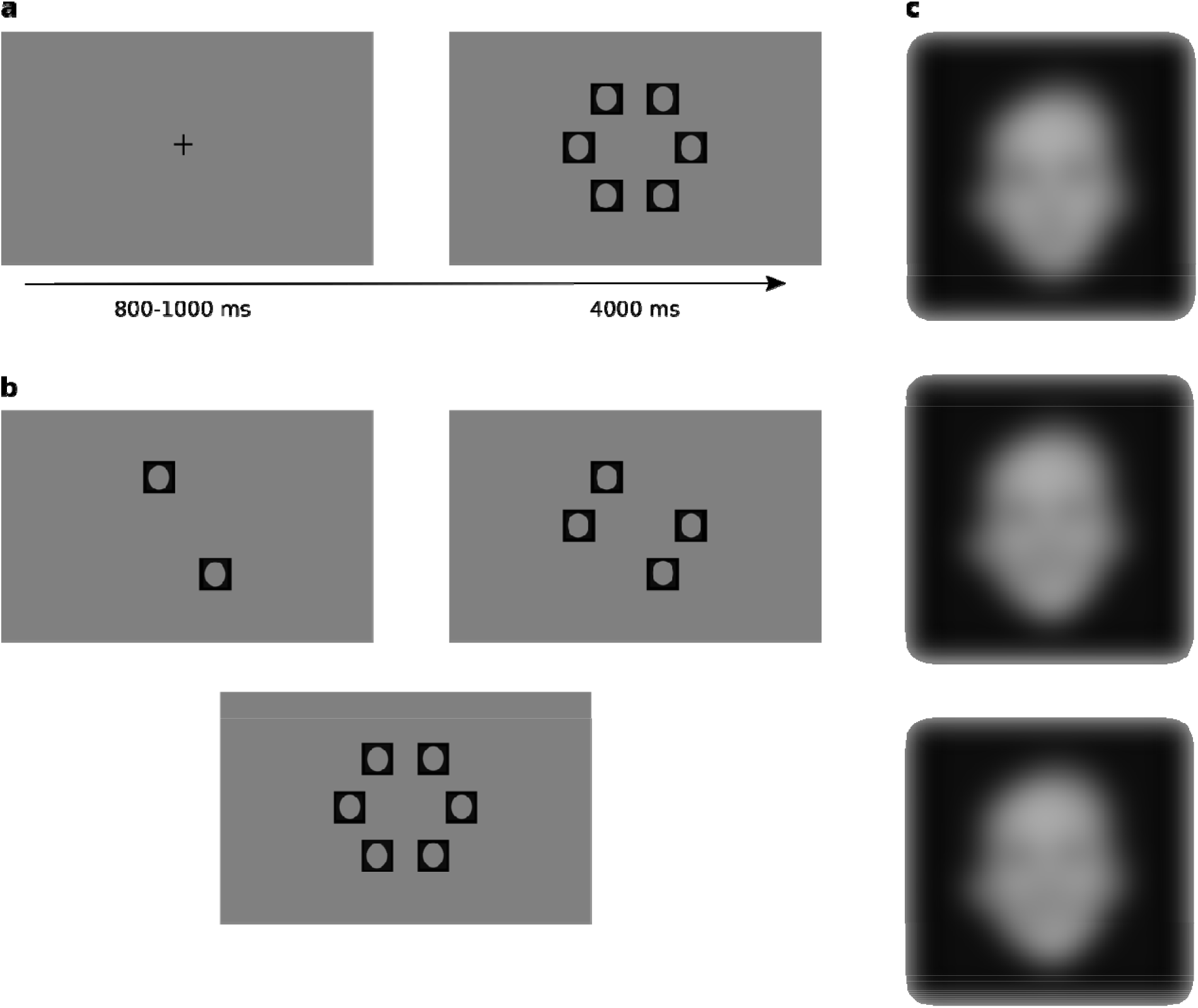
a) Example sequence for a single target present trial for the happy expression. The fixation cross was on screen for a jittered interval between 800 and 1000 ms. The array of face stimuli stayed on screen until the participant responded, or for a maximum of 4 s. The subsequent trial began immediately after the participant’s response. b) Example stimulus arrays for sets of 2, 4 or 6 images. c) Example familiar target with neutral, happy and angry expression. [This figure is blurred for the preprint]

### Experimental setup

The experiment was run on a GNU/Linux workstation with presentation scripts written in MATLAB (R2014b) using Psychtoolbox (version 3.0.12). The resolution of the screen was 1440 × 900 pixels and the refresh rate was 60 Hz. Participants sat approximately 50 cm from the screen with their face resting on a chin rest. All stimuli were presented against a uniform gray background.

### Task

The first phase of the experiment involved familiarization with the target stimuli. The participants saw each image of familiar and unfamiliar targets making angry and happy expressions twice, for 4 seconds each time, with 500 milliseconds interstimulus interval between each image.

Following the familiarization phase, participants performed 10 practice trials identical in structure to the experiment (the practice trials were not included in the final analysis). After the practice trials, the experiment started and participants performed 2 blocks of 192 trials each. In one block, the target was a face with an angry expression among neutral distractors. In the other block, the target was a face with a happy expression among neutral distractors. The identity of the target stimulus with an angry or happy expression was familiar on half of the trials and unfamiliar in the other half. On target present trials, the identities of distractor faces with neutral expressions were always unfamiliar. On target absent trials, one face with neutral expression was a familiar identity on half of the trials. The order of blocks was counterbalanced across participants. Participants were asked to press the left arrow key if the target was present and the right arrow key if the target was absent. They had a maximum of 4 seconds to make their response. Participants were instructed to respond only to the expression of the target and to ignore the identity of the target. Participants were also instructed to respond as quickly as possible, but not at the expense of accuracy.

Each block began with the instructions specifying which expression was the target for the following trials (e.g. “look for happy expression”). Each trial ended when the participant’s response was recorded or ended after a maximum of 4 seconds. The intertrial interval was jittered between 800 milliseconds and 1 second. Participants fixated a central fixation cross during the inter-trial interval, and the cross disappeared once the face stimuli appeared, allowing eye movements to the stimuli. Every 48 trials, the participant was given the option of taking a short break. For each block, each unique trial type was repeated 4 times. A trial type is defined by a given set size (2, 4 or 6), target condition (present or absent) and the unique identity of the target (8 identities — 4 familiar, 4 unfamiliar). The sex of the distractors always matched the sex of the target. Targets with the emotional expression were presented in half the trials for each block. Targets were equally presented in the left and right hemifield.

After the visual search session was over, participants rated the target stimuli on how recognizable the happy and angry expressions were. In order to rate the expressions of the targets, participants judged how clearly the face expressed the emotion of anger or happiness by choosing a number between 1 and 5 (with 5 being maximally expressive).

### Statistical analyses

Accuracies were analyzed using the function *glmer* from the package *lme4* (Bates et al., 2014) and the function *Anova* from the package *car* (Fox et al., 2012). The analysis of accuracy involved fitting a generalized linear mixed model to the data — with accuracy as the dependent variable; target presence, set size, target expression, target familiarity and the interactions between these variables as fixed effects; the participant and stimulus identities as random effects. We also included the sex of the target as a fixed effect in our model. We compared models with different random slopes and intercepts until we identified the best model with the lowest Akaike’s Information Criteria (AIC). To identify which fixed effects significantly affected the accuracy, we used Type 3 analysis of deviance (using Wald’s □^2^ test).

For the analysis of reaction times, we used the function *lmer* from the package *lme4* (Bates et al., 2014) and the function *Anova* from the package *car* (Fox et al., 2012). We fitted our data with a logit mixed model with log transformed reaction times as the dependent variable and target presence, set size, target expression, target familiarity and the interactions between these variables as fixed effects. We also added the sex of the target as a fixed effect. The random effects included participants and stimulus identities. The model complexity was reduced by removing random slopes and intercepts until the best model was identified using the lowest AIC value. Again, we used Type 3 analysis to determine the significance of the fixed effects entered in the model. The bootstrapped confidence intervals for the figures and the effect sizes were estimated with custom code written in R. The 95% confidence intervals were calculated by randomly sampling subsets of trials from each participant across conditions to compute the dependent variable (accuracy, reaction time and rating). For both models, we used a polynomial contrast for set size and zero sum contrasts for the remaining fixed effects.

## Results

### Accuracy

We fitted generalized linear mixed models to our data, with accuracy as the dependent variable. We used two separate models to analyze the accuracy for the target present and absent conditions, consistent with prior literature (Tong & Nakayama, 1999). For the target absent trials, participants performed at nearly 100% accuracy across all conditions (Figure 2). For detail, see Supplemental Material.

**Figure 2:**
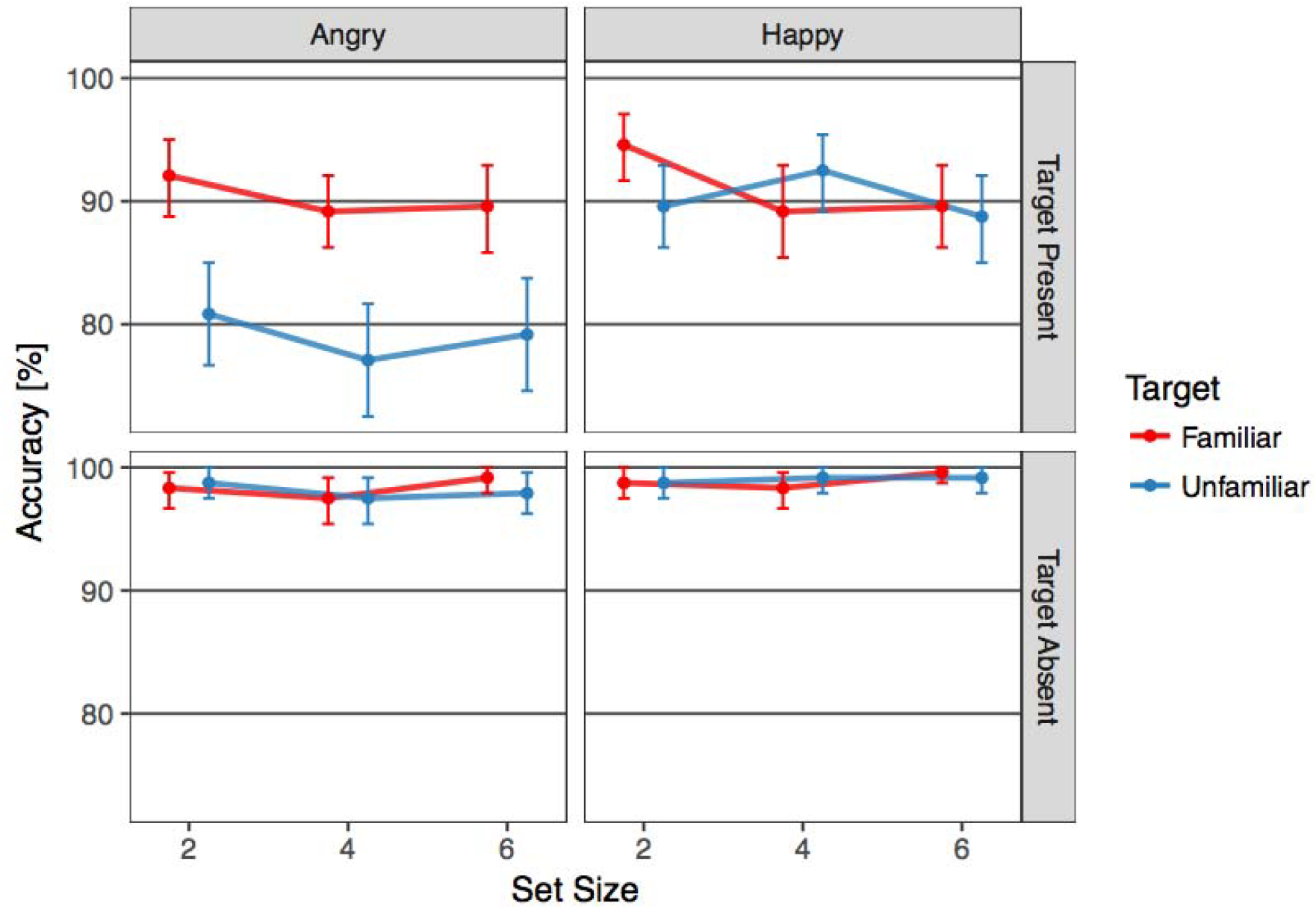
Accuracy in reporting presence or absence of target expression as a function of set size of images. Rows depict target present or absent condition and columns depict the target expression for the block. The effect of familiarity on accuracy is driven by greater accuracy for familiar faces when the target is present and the target expression is angry. Error bars indicate 95% bootstrapped confidence intervals.

In the target present condition, the fixed effects in the model with the lowest AIC were set size, familiarity of the target, target expression and their interactions. We also added the fixed effect of target sex. The random effects for this model included slopes and intercepts for participants. The set size of the items in the search array did not have a significant effect on the accuracy (*2*(*2*) = *4*.*74*, = *0*.*09*). We found significant effects of familiarity (2(1) = 21.03, LJ < 0.001) and expression (*2*(*1*) = *21*.*43*, □ < *0*.*001*). The interaction between these two fixed effects was also found to be significant (*2*(*1*) = *12*.*15*, □ < *0*.*001*) while none of the other interactions were found to be significant (set size x familiarity: (*2*(*2*) = *4*.*41*, □ = *0*.*1*), set size x expression:( *2*(*2*) = *1*.*16*, □ = *0*.*55*), set size x familiarity x expression: (*2*(*2*) = *3*.*0*, □ = *0*.*22*)). Lastly, we also found a significant effect of the sex of the target (*2*(*1*) = *4*.*02*, □ = *0*.*04*). (Figure 2) The analysis of individual contrasts revealed that subjects were more accurate for familiar targets overall (Familiar: 94.65% [93.96,95.35], Unfamiliar: 91.60% [90.73, 92.43]), and for targets with a happy expression as compared to the angry conditions (Happy: 94.83% [94.10, 95.56], Angry: 91.42% [90.56, 92.29]). The significant interaction between the two terms reflected that the advantage of familiarity on accuracy in the target present condition was found only for angry targets (90.28% [88.33, 92.08] versus 79.03% [76.53, 81.53]) and was absent for happy targets (91.11% [89.17, 92.92] versus 90.28% [88.33, 92.22]).

### Reaction Times

Only correct trials were included in the analysis of reaction times. We fitted a linear mixed model to our data, with log normalized reaction times as our dependent variable. As with accuracy, we analyzed reaction times in the target present and absent conditions separately. The main effects in the model with the lowest AIC included set size, familiarity of the target, target expression and the interactions between these variables. We also included the main effect of target sex in the model. The random effects included slopes and intercepts for the participants and the slopes for the different image combinations. We found a significant effect of set size, target familiarity, target expression and target sex. The interaction between target familiarity and expression was also found to be significant, while none of the other interaction terms were significant (set size x familiarity:, set size x expression:, set size x familiarity x expression:). Overall, in the target present condition, familiar targets were detected 135 ms [95,175] faster than the unfamiliar targets for angry expressions, and 1 ms [-33,31] slower for happy expressions (Figure 3).

**Figure 3:**
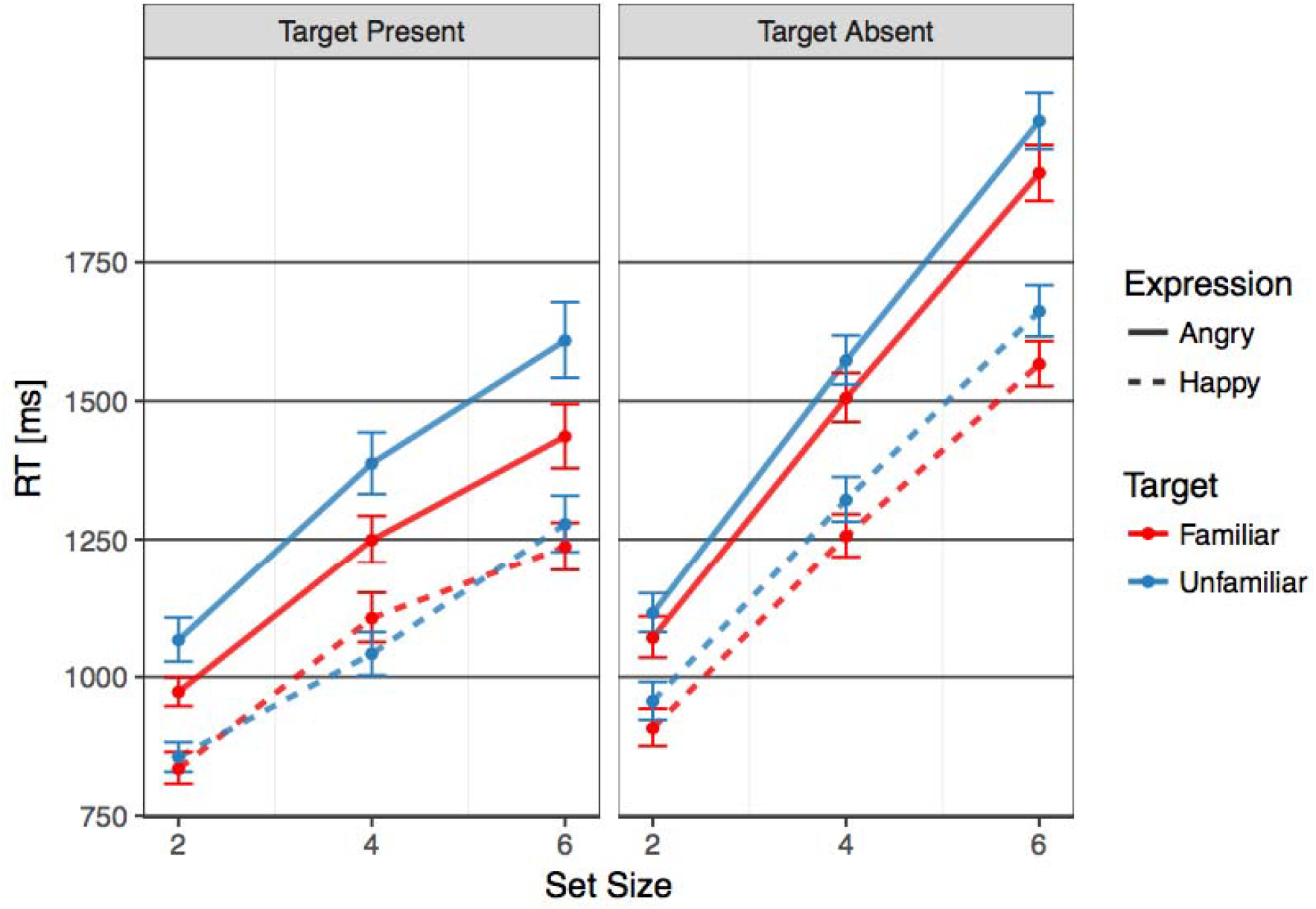
Reaction times (ms) as a function of set size of images. Longer reaction times were observed for unfamiliar faces when the target was an angry expression. Error bars indicate 95% bootstrapped confidence intervals.

In the target absent condition, the dependent variable was log transformed reaction times and the main effects were set size, target familiarity, target expression and the interactions between these terms. Target sex was also included as a fixed effect. The random effects included random slopes and intercepts for participants and random slopes for different image combinations. We found significant main effects of set size (*2*(*2*) = *1964*, □ < *0*.*001*), target familiarity (*2*(*1*) = *18*.*71*, □ < *0*.*001*), target expression (*2*(*1*) = *449*.*53*, □ < *0*.*001*), and target sex (*2*(*1*) = *4.67*, □ = *0*.*03*). None of the interactions were found to be significant (familiarity x expression: (*2*(*2*) = *0*.*07*, □ = *0*.*78*), set size x familiarity: (*2*(*2*) = *0*.*29*, □ = *0*.*86*), set size x expression:( *2*(*2*) = *3*.*46*, □ = *0*.*17*), set size x familiarity x expression: (*2*(*2*) = *0*.*0*, □ = *0*.*99*)). Interestingly, despite the absence of a target making an expression, we found that subjects were faster in making a response when the task was to report the presence of a target with a happy expression (1.28 s [1.26,1.29]) as compared to the angry expression (1.53 s [1.51,1.55]) (Figure 3). Moreover, even though no targets with expressions were present in the stimulus array — either a familiar or an unfamiliar identity with a neutral expression was included as a distractor –– it was, therefore, possible for us to analyze the effect of the presence of a familiar distractor in the search array even in the trials when there was no target with an emotional expression. We found that participants responded that a target was absent faster when a familiar face with a neutral expression was among the stimuli, as compared to target absent trials with all unfamiliar faces. This difference was seen both on angry target and happy target trials (69 ms [33, 105] and 70 ms [38, 101] differences for angry target and happy target trials, respectively). Unstandardized effects for reaction times in both target present and absent conditions are depicted in Figure 4.

**Figure 4:**
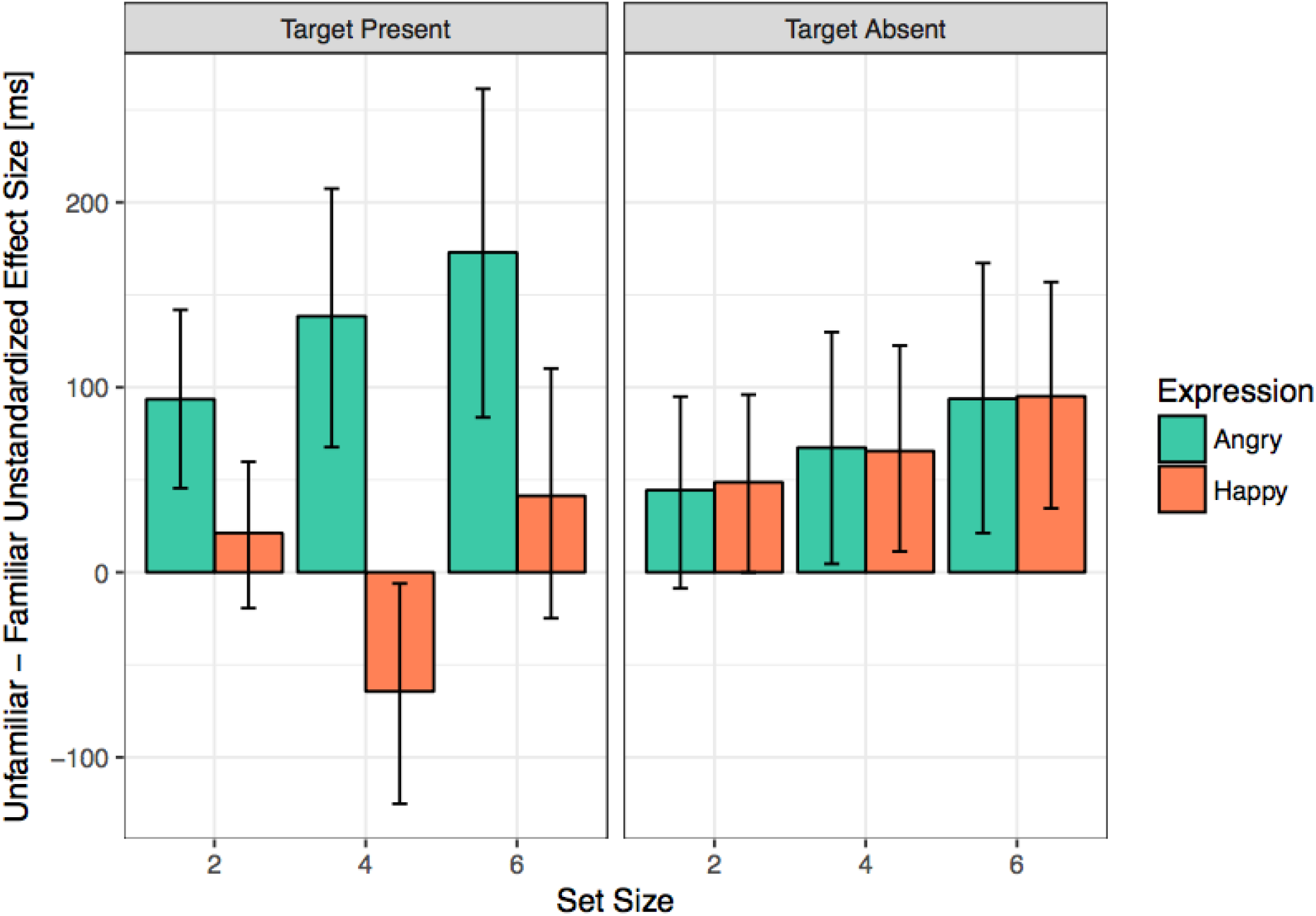
Unstandardized effect sizes for the difference in reaction times between unfamiliar and familiar faces as a function of set size. Error bars indicate 95% bootstrapped confidence intervals.

### Expression Ratings

We also analyzed the ratings for how recognizable happy and angry expressions were in each target identity. We fitted a generalized linear mixed model to the rating data, with the rating for how recognizable the expression was as the dependent variable, and target familiarity, expression and sex as independent variables. The model with the lowest AIC included subject specific slopes and intercepts as random effects. We used the poisson distribution as the linking function for this model.

We found a significant effect of target sex on the ratings, which was driven by lower recognizability ratings for the males (3.91 [3.83,3.99]) as compared to the females (4.53 [4.46,4.59]). We did not find a significant effect of the target expression or of target familiarity. The interaction between target familiarity and expression was not significant, nor was the trend for ratings of familiar versus unfamiliar angry expressions (Figure 5).

**Figure 5:**
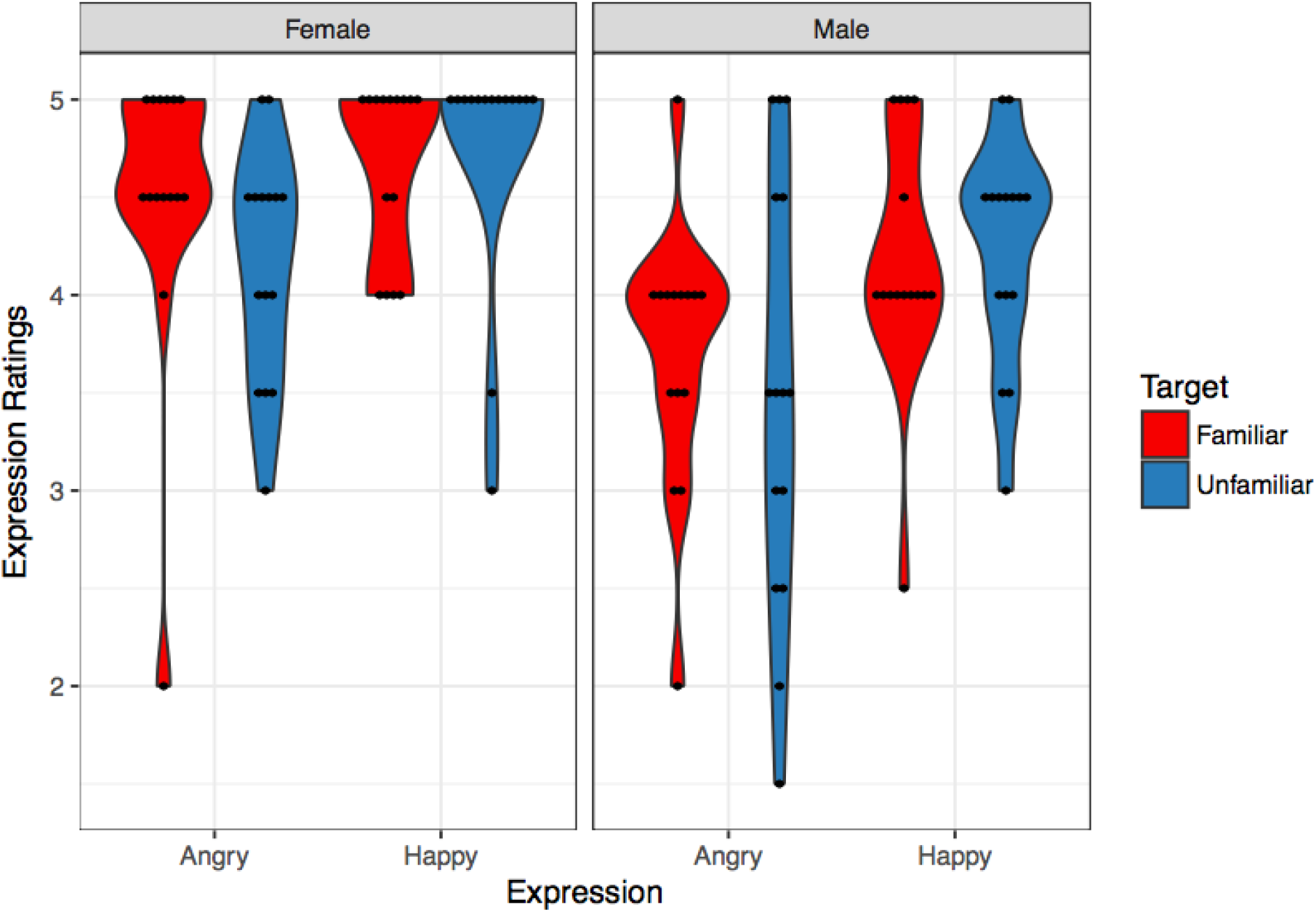
Ratings for recognizability of expression (1: not expressive, 5: very expressive) as a function of target expression. Left panel is for female targets, right panel is for male targets. Ratings were lower for male targets as compared to female targets and angry expression compared to happy expression. Error bars indicate 95% bootstrapped confidence intervals.

## Discussion

In this study, we investigated whether personal familiarity facilitates the detection of emotional facial expressions. The results confirmed our hypothesis for detection of angry facial expression but not, interestingly, for happy expressions. We found that participants were more accurate and faster at detecting an angry facial expression if the face making that expression was personally familiar. The effect was large, with an 11% difference in detection accuracies and a 135 ms advantage in detection time. With unlimited time, the angry expressions of unfamiliar faces were rated as being equally expressive as the angry expressions of personally familiar faces, indicating that the accuracy and speed differences in the visual search task were not due to differences in expression ambiguity.

Previous research has shown that familiar identities are detected faster, as compared to unfamiliar identities (Gobbini et al. 2013; Visconti di Oleggio Castello and Gobbini, 2015; Ramon et al. 2015; Visconti di Oleggio Castello et al. 2017). Moreover, the effect of familiarity on detecting familiar identities is robust when face images are inverted (Visconti di Oleggio Castello et al. 2017), suggesting that familiarity-based facilitation can also involve parts-based, rather than holistic, face perception processes. Social cues, such as head angle and eye gaze, also are detected faster when conveyed by familiar faces, as compared to unfamiliar faces (Visconti di Oleggio Castello, Guntupalli, et al., 2014), with a speed advantage similar to that found in the current study for detection of angry facial expressions.

The current study shows that familiarity-based facilitation of social cue detection extends to emotional expressions, which are conveyed by contractions of facial muscles (Ekman, 1993;Martinez et al., 2012). Thus, the facilitation of visual processing of faces that accrues from learning socially-salient, personally familiar faces involves both the detection of invariant features that specify identity and the detection of changeable features that convey facial gestures and social signals – processes that are mediated by the ventral and dorsal pathways, respectively, in the core system of the human neural system for face perception (Haxby et al. 2000; Haxby & Gobbini, 2011; Visconti di Oleggio Castello, Halchenko, et al. 2017). We have shown further that visual learning of familiar faces affects retinotopic biases for identity recognition, suggesting that this visual learning extends to early processes in retinotopic visual cortex (Visconti di Oleggio Castello et al. 2018). This body of work is consistent with our hypothesis that visual learning of familiar faces involves the development of detectors for individual-specific fragments of a familiar face that facilitate detection of that face’s identity and gesture.

Familiar face perception also spontaneously evokes representations of person knowledge — that person’s dispositions, personality, and position in a social network (Gobbini et al. 2004; Leibenluft, et al. 2004; Gobbini & Haxby, 2007; Arsalidou et al., 2010; Taylor et al., 2009; Natu & O’Toole, 2011; Sugiura et al., 2014; Visconti di Oleggio Castello, Halchenko et al., 2017; Ramon & Gobbini, 2018). The effect of person knowledge on detection of facial expression, which may be mediated by a top-down mechanism, may also play a role in familiarity-based facilitation.

Interestingly, familiarity-based facilitation did not extend to detection of happy expressions. Participants detected happy expressions conveyed by familiar and unfamiliar faces with equivalent accuracies and at equivalent speeds. Our stimuli with happy expressions were detected faster and more accurately than were our stimuli with angry expressions. In general, others have found that happy expressions are recognized with highest accuracy compared to the other canonical expressions (Kirouac & Doré, 1983; Kirita & Endo, 1995; Leppänen et al., 2003; Wells et al., 2016). The absence of an effect of familiarity on the detection of the happy expression suggests that this process is optimized, perhaps because it is such a common social cue for interactions with both familiar and unfamiliar others. This effect may also be explained by low-level features that distinguish happy and neutral expressions such as exposed teeth, even though we did not observe a pop out effect in the visual search task. Further research could determine if this familiarity invariance extends to subtler, unposed, or less stereotypic expressions of happiness. Similarly, further research could determine whether familiarity-based facilitation is also found for other standard expressions, such as disgust and surprise, as well as for non-standard expressions, such as contempt, boredom, and skepticism.

We found an unexpected effect of familiarity on target absent trials. Faster responses on target absent trials in which one stimulus was a familiar face with a neutral expression suggests that participants adopted a strategy in which they may have terminated search when they detected this familiar distractor, forgoing examination of the other unfamiliar distractors. This suggests that the participants were using the strategy of performing a self-terminating visual search (Sternberg, 1966), and relying on the familiar face as an indicator of whether an emotional expression target was present or absent.

The face perception system plays a central role in social interactions and mounting evidence shows that it is optimized for interactions with personally familiar others. This optimization is evident in both detection of invariant facial features that specify identity and changeable facial features that carry social signals. Optimization involves visual learning, based on extended naturalistic interactions with personally familiar others, and involves both the ventral and dorsal face pathways in the distributed system for face perception and extends to early processes in retinotopic cortex. Familiar face perception, unlike unfamiliar face perception, also involves the spontaneous activation of neural systems for the retrieval of person knowledge. Understanding face perception – its perceptual and cognitive processes, its neural substrates – requires understanding how it processes familiar faces, in much the same way that understanding language processing and its neural substrates requires understanding how one processes one’s native language.

